# Evidence for subjective values guiding cognitive integration of gravity in a free speed manifold pointing task

**DOI:** 10.1101/2023.01.23.525134

**Authors:** Arthur Devemy, Charalambos Papaxanthis, Lucieny da Silva Pontes, Thierry Pozzo, Pauline M. Hilt

## Abstract

Studies of regularities in human movement control have led to the idea that our brains incorporate generic motor laws that guide our behaviors and thus these regularities. Among these, it has been argued that a generic internal model of gravity minimizes the body’s energy expenditure during actions performed in the vertical plane. According to this hypothesis, the energy criterion would represent the factor dominating motor planning. However, a growing body of experimental evidence indicates the presence of significant individual sensorimotor differences that suggest the use of various motor decision criteria.

We asked 41 voluntary participants to perform a free speed manifold pointing task (i.e., at three free different speeds and toward non-salient targets), in the vertical plane designed to avoid situation-specific responses and favor the expression of singularities. We found that temporal parameters (duration of motion and vigor) were idiosyncratic and not affected by mechanical effects induced by changes in velocity and gravitational force (i.e., assistive or resistive forces). Upward motions had a shorter time to reach maximum velocity than downward motions confirming the presence of asymmetric movement timing. Finally, synchronization of up-down movements was not correlated with energy minimization, indicating that movements performed in the vertical plane do not rely exclusively on a standard representation of gravity reducing effort. Rather, we concluded that individual values and the subjective rewards associated with each of these values could influence the planning of movements executed along the vertical plane as well as the cognitive integration of the gravitational force field.

## Introduction

The presence of motor regularities recorded during human movements performed in different contexts is recurrently evoked as the expression of generic representations of actions. Fitts’ law (Fitts, 1954) and the 2/3 power law (Grasso et al., 2002; Lacquaniti et al., 1983; Viviani & Flash, 1995) are examples of kinematic regularity extraction suggesting the existence of common sensorimotor representations anticipating mechanical consequences due to body-environment interactions. Following this theoretical framework, it has been proposed that the mechanical effect of the gravitational force field on the whole body is anticipated via a common representation generating adequate sensorimotor interaction in 3D space (Berret, Darlot, et al., 2008; Papaxanthis et al., 1998). In other words, an internal model would minimize the effort by taking advantage of the gravitational force. Specifically, kinematic characteristics that vary with the direction of motion (i.e., upward motions have a shorter time to reach maximum velocity than downward motions, see (Papaxanthis et al., 1998) would reflect the best motor solution (Gaveau et al., 2021). In this view, the mechanical effect of the gravitational force would thus be anticipated in a uniform way, via the estimation of the energy cost of the actions carried out with or against the gravitational force field (Gaveau et al., 2016). Walking or running during endurance events, or when using robotic exoskeletons that create new energy context (Selinger et al., 2019) provide many examples supporting the principle of optimizing energy expenditure. Yet, during short-duration movements, the central nervous system (CNS) is less likely to deplete energy stores and the minimization of energy costs probably matters less in the decision process that guides the type of response chosen by the subjects. In many cases, humans will choose an action requiring greater effort, despite the availability of a lower-effort option, such as when fleeing from danger or when thirsty they quickly grab a glass of water. Similarly, when reaching an object in a force field where a straight path requires more force than a curved path, subjects have been shown to consistently choose the straight path (Huang et al., 2012; Izawa et al., 2008; Kistemaker et al., 2010; Shadmehr et al., 2016). All these observations showed that minimizing energy expenditure via a stereotyped common neural representation may not be the only concern of the nervous system during action preparation. Moreover, a simple motor solution maximizing energy saving assumes a subject-independent representation of gravitational force. However, several recent articles have highlighted the role of individual internal values (this term refers here to the subjective process guiding the motor response) in motivating the participant to choose one action over another (Hilt et al., 2016, 2020; Labaune et al., 2020; Shadmehr et al., 2016; Słowiński et al., 2016). During a whole body pointing task, the presence of discrete individual postural strategies highlighted the role of two internal values to minimize the mechanical energy, or to maximize the body equilibrium (Hilt et al., 2016).

This study aims to verify the potential role of subjective values in the planning of arm-pointing movements performed with or against the force of gravity. Although interesting, previous approaches have mainly focused on the idea that gravity is integrated as a force to be compensated (Atkeson & Hollerbach, 1985; Flanders & Herrmann, 1992; Guigon et al., 2007; Hollerbach & Flash, 1981; Olesh et al., 2017) or to anticipate (Gaveau et al., 2014, 2016; Gaveau & Papaxanthis, 2011; Gentili et al., 2007; Le Seac’h & McIntyre, 2007) in order to ultimately minimize the effort required. However, reducing the control of a task to its energy optimization confines the understanding of the cognitive integration of the gravitational force to the single criterion of a homeostatic balance of the biological system, while other motivations can influence the realization of the movement. More important, the internal representation of gravity was exclusively investigated under restricted conditions that reduce motor variability by forcing the subject to reach a salient target as quickly as possible. Alternatively, when the decision-making process guiding the kinematic options is subject to less strict experimental constraints, we believe that the cost optimization criterion could become blurred, so that other optimizing mechanisms would be revealed.

A pointing task performed at maximum speed towards a salient target probably homogenizes individual responses to a generic behavior that results from biomechanical limitations thereby masking potential inter-subject variability. Conversely, by reducing the specificity of the protocol and the boundary condition of experimental context, the expression of the singularities previously masked could be facilitated. In order to avoid excessive normalization of the data, which also limits the generalizability of the results (Simons et al., 2017) and offers more freedom to the subject, we designed a protocol that minimizes spatial (target redundancy) and temporal (spontaneous movement speed) constraints (Berret et al., 2011). On the one hand, the consistency of some task-relevant parameters whatever the direction and the speed of the movement would confirm the primary role a simple effort cost solution adopted by all subjects. On the other hand, the inconsistency of some kinematic parameters in addition to significant inter-individual differences would indicate that the minimization of mechanical energy expenditure alone cannot account for this wide range of behaviors.

## Methods

### Participants

Forty-one healthy right-handed participants (15 males and 26 females, mean age: 21), with normal or corrected-to-normal vision and naïve to the task, participated in this study. None of the participants had neurological, psychiatric, or other relevant medical problems affecting motor performance. Each participant provided written informed consent before participating in the study. The Ethics Committee of the Université de Bourgogne approved the protocol.

### Procedure

The participants were seated in a comfortable armchair in a quiet darkened room. An LCD computer monitor (1920×980 pixels; refresh rate 60 Hz) was placed at a distance of 90 cm from their frontal plane. Participants were required to perform a free-speed manifold pointing task in the vertical plane. They had to move their right arm from an initial red cross to a final memorized blue area (circle 3 cm in diameter). All targets were centered on the participant’s right shoulder at a distance equal to the length of their fully extended arm. Two movement directions (two levels: Downward and Upward, further referenced as Down and Up) and three execution speeds (three levels: Slow, Spontaneous, and Fast, further referenced as Fast, Spont, and Slow respectively) were tested.

Precisely, each trial started with a red cross appearing at the top (Down condition) or bottom (Up condition) of the monitor (Figure 1). The participant was required to point at this initial red cross and remain there. Once he adopted this position, the red cross disappeared, and a blue circular area appeared (for 0.5 s) in the opposite location on the vertical axis of the monitor. After its disappearance, the participant had to move his arm from the initial position to one own-selected location within the memorized final zone. The use of a final area (rather than a salient target) allows the final position of the finger to be unconstrained and frees up the participant’s movement options, thus creating a decision context (Berret et al., 2011). The delay between the disappearance of the red cross and the appearance of the blue area varied between 0 s, 1 s, and 2 s to communicate the speed condition. These delays are intended to suggest to the participant the realization of a particular speed condition (0 s to evoke Fast, 1 s to evoke Spontaneous, and 2 s to evoke Slow).

**Figure 1.**
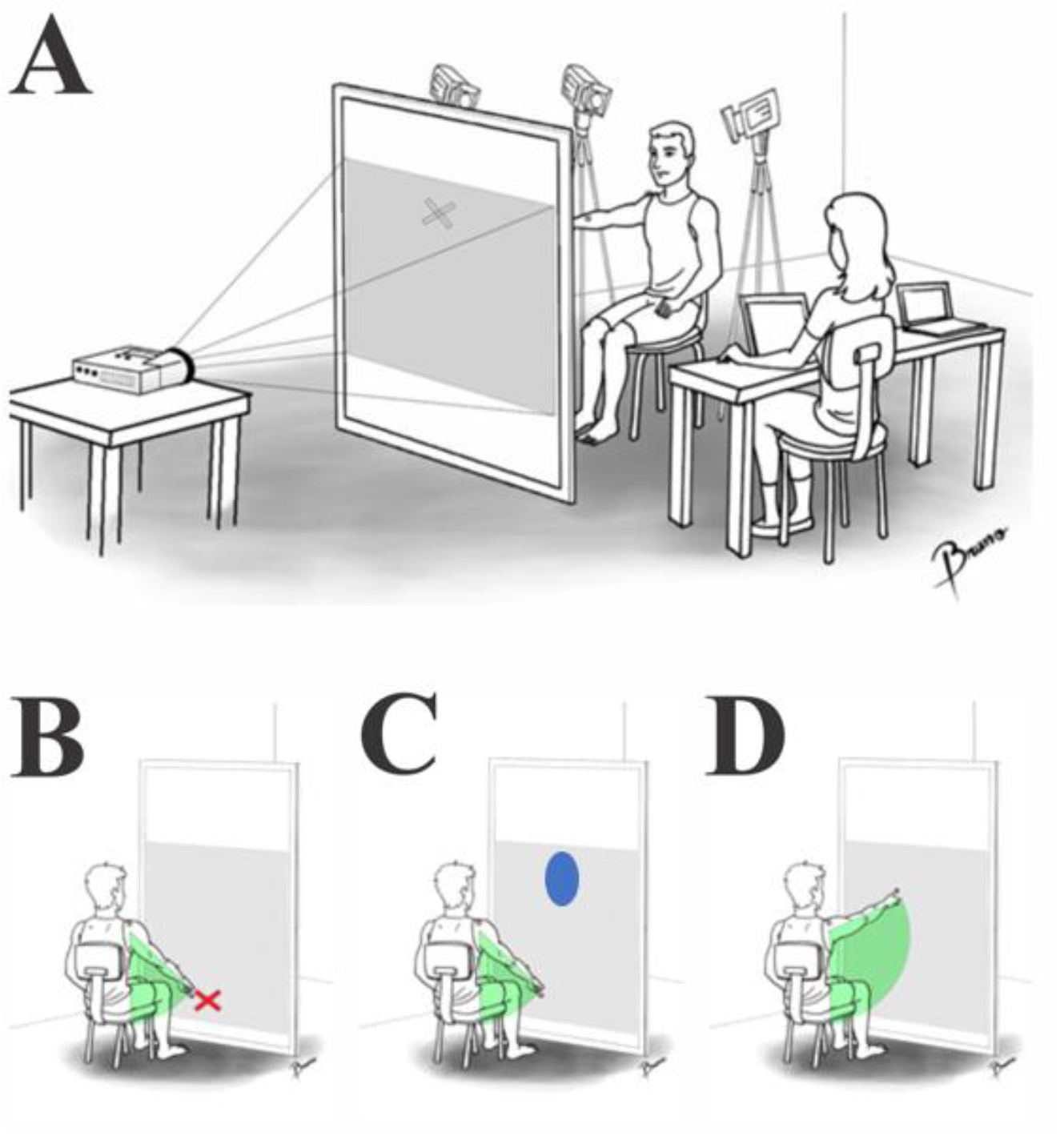
Illustration of the experimental setup. A) The participant was seating in a quiet darkened room, in front of a LCD screen. B) Each trial started with the appearance of an initial red cross. C) After a variable delay indicating movement speed, a final blue area appeared at the vertical opposing location of the screen D) After disappearance of the blue area, the participant performed the vertical movement from the red cross to the blue area location.

Each participant executed 20 trials per condition, resulting in 120 trials in a completely randomized order. Each trial lasted for approximately 5 s, resulting in a total of approximately 25 min of recording. Every 20 trials, a 2-minutes pause was proposed to the participant.

We performed a calibration phase before the experiment and after each pause to accurately estimate the perceived locations of the initial and final targets for each subject. In these phases, the participants were asked to successively point at the top and bottom crosses steadily while being recorded. For each subject, these two positions were used as references to compute the constant and variable errors of reaching movements.

Finally, to define the subjective level of speed expected in the speed condition, each participant attended a training phase (before the experiment). During this phase, participants were first asked to perform pointing movements in the vertical plane at their own selected preferred speed (Spont condition). Indeed, the Spont condition was designed to assess the behavior of the participants when they were able to choose the speed of realization themselves, and thus select the motor strategy that seemed most appropriate to them. In the training phase, we next asked them to still perform pointing movements in the vertical plane, but slowing down (Slow condition) or speeding up (Fast condition) their movements. They performed three trials in each direction (Up and Down) for each speed level (i.e., a total of 18 trials). These self-defined speeds then corresponded to the speed levels required of that participant in each experimental speed condition of the main experiment. The Fast and Slow conditions used here aimed at testing the influence of a speed constraint on the choice of a motor strategy in this type of pointing task. For example, when performing movements at abnormally slow speeds, movement replanning may induce inconsistencies between the trials of the same participant. Inversely, a speed that is too fast can homogenize individual data (reduce intra- and inter-subject variability) and introduce a standard and uniform way of moving, thus masking the existence of individual motor signatures.

### Materials

Reaching movements in three axes (mediolateral, X; antero-posterior, Y; and vertical, Z) were recorded using a Vicon Motion Capture setup (Vicon, Oxford, UK) with 3 cameras recording at a frequency of 100 Hz. Participants were equipped with four motion captors (15 mm in diameter) placed at the following anatomical locations: the acromial process (named here “shoulder”), the lateral condyle of the humerus (named here “elbow”), the styloid process of the ulnar (named here “wrist”), and the last phalanx of the index finger (named here “index”). The presentation of the stimuli was controlled using Psychtoolbox Version 3.0 (PTB-3), implemented in MATLAB (The MathWorks Inc.).

### Motion analyses

All analyses were performed using custom software written in Matlab (MathWorks, Natick, MA, USA). The 3D coordinates of all markers were low-pass filtered (4.0 Hz cut-off 4th order Butterworth filter (Scholz & Schöner, 1999). Movement onset and offset were defined as the moment at which the velocity of each subject’s index finger reached or fell below 5% of the peak velocity for each trial (Hilt et al., 2016). All analyses of hand movement variables were performed over the period between these onsets and offsets.

To assess the consistency of pointing movements in the spatial domain, we calculated the Constant Error (CE, in mm), defined as the vertical (Z axis) distance between the movement offset position and the center of the final area (recorded during the calibration phase), and the Variable Error (VE, in mm), defined as the Standard Deviation of the distance between the movement offset position and the center of the final area.

To evaluate the temporality of the movement, we computed three standard kinematic parameters: Movement Duration (MD, in seconds), defined as the duration between movement onset and offset; Maximum Velocity (Max V, in mm.s-1), defined as the peak vertical (Z axis) velocity of each trial; and Time to Peak Velocity (TPV), defined as the ratio between Acceleration Duration (i.e., from the onset time to the Peak Velocity time) and MD. To investigate the intra-subject variability of these variables, we computed the coefficient of variation (CV) of MD and TPV, defined as the ratio between the inter-trial Standard Deviation and the mean of each variable.

Furthermore, to obtain a direct measure of the temporal changes potentially associated with movement direction, we computed the TPV delta (Δ_Down-Up_) at each trial level, defined as the difference in TPV values between the Downward and Upward directions.

Additionally, to evaluate the interindividual consistency of speed-related variables, we computed individual vigor scores relating individual MaxV to the general distribution of MaxV obtained across the 41 subjects. Individual vigor scores were computed according to the following formula used by (Labaune et al., 2020):

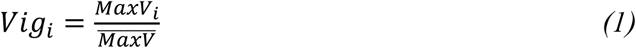

where *Vig_i_* is the Vigor score of participant *i, MaxV_i_* is the mean Max V of participant *i* and 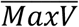 is the mean Max V across all participants.

Finally, to assess how participants’ behavior may be related to the minimization of a certain energy criterion, we computed the rate of metabolic energy expenditure for each subject. We used the formula from (Shadmehr et al., 2016)designed for horizontal reaching movements and adapted it to vertical arm movements by adding the weight of the work of gravity torque as follows:

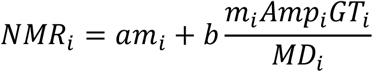

Where *NMR_i_* is the mean net metabolic rate for subject *i, m_i_* is the estimated arm weight (2.71 % of subject body weight), *Amp_i_* is subject’s movement Amplitude, *MD_i_* is subject’s movement duration, *GT_i_* is the work of gravity torque of subject *i*. *a* and *b* were used as weights and fixed so as to get values comparable to (Shadmehr et al., 2016; a=15 and b=0.1).

The Work of Gravity Torque of each subject was computed using the following equation:

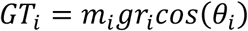

where *GT_i_* is the work of gravity torque for subject *i, m_i_* is the estimated subject’s arm weight (2.71 % of body weight), g is the earth constant gravitational acceleration, r_i_ is subject’s arm length, and θ_i_ is the angle between the arm orientation and the vertical axis.

### Statistical analyses

All statistical analyses were performed using R software (Version 4.1.2). A repeated-measures two-way analysis of variance (ANOVA) was performed on each dependent variable (MD, Vmax, CE, VE, Amp, and TPV), with speed (Spontaneous, Fast, Slow) and direction (Down, Up) as within-subject factors. Tukey’s HSD test was used as a post-hoc test when needed (α=0.05). Correlations were determined using Pearson’s correlation test. P-values were adjusted for multiple comparisons using the False Discovery Rate (FDR) method (p.adjust function in R).

## Results

### Movement Duration

The 3×2 ANOVA ran on Movement Duration, showed a main effect of speed (F(2, 39)=139.09, p<0.0001), but no main effect of movement direction (F(1, 40)=2.1, p=0.15) nor interaction (F(2, 39)=0.30, p=0.73). This confirms that all participants were able to perform the task at three different speeds, as required by the protocol. In particular, MD significantly increased (71%, p<0.001) and decreased (41%, p<0.001) in the slow and fast conditions, respectively, compared to the spontaneous (Spont) speed (Table S1, Supplementary Materials). MD was strongly subject-dependent ranging from 0.68 to 3.75 sec (1.70±0.68 s; MEAN±STD), 0.61 to 1.73 sec (0.99±0.26 s), and 0.26 to 1.17 sec (0.58±0.17 s) in the Slow, Spont, and Fast conditions respectively (Figure 2A).

**Figure 2.**
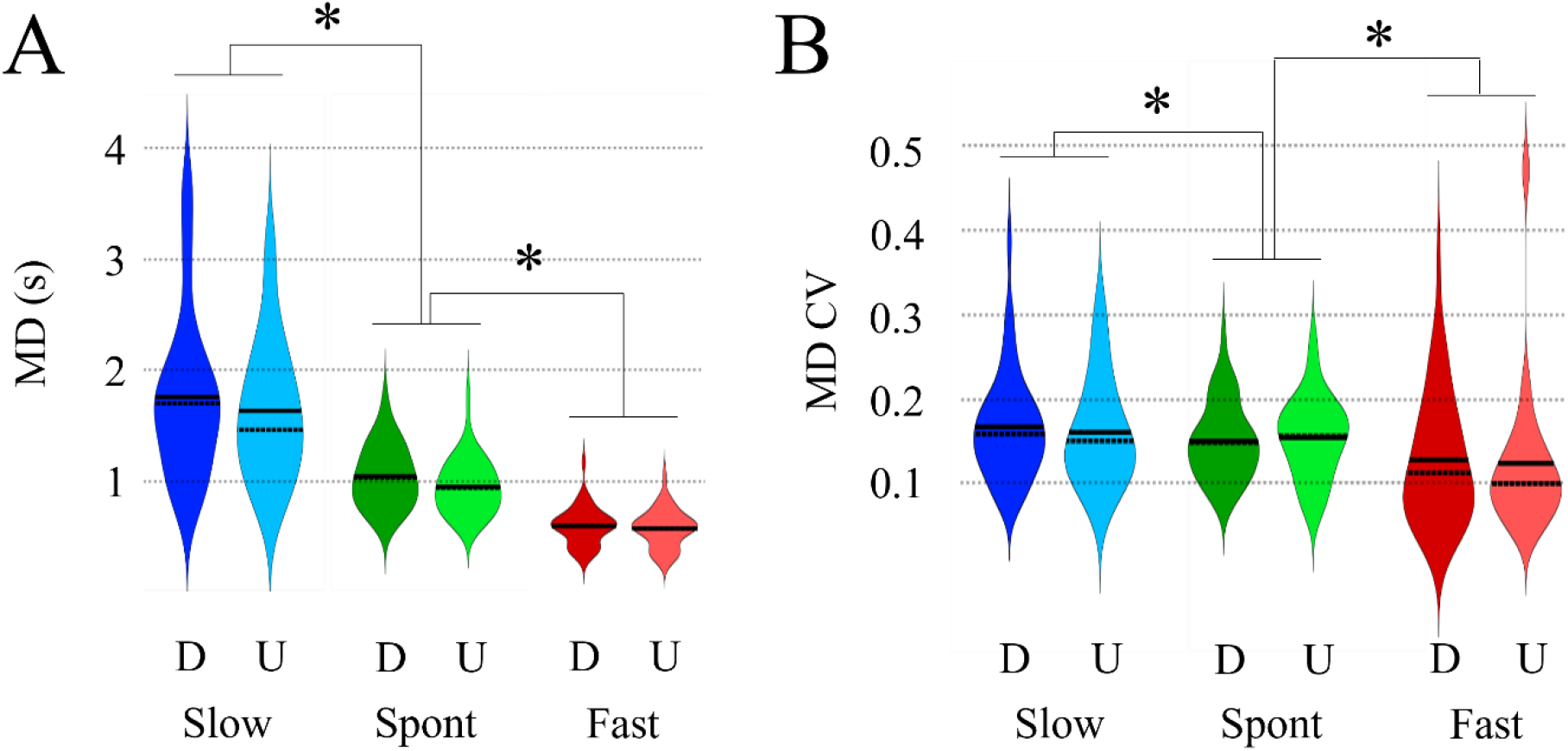
Effect of direction and speed on movement duration (MD; in seconds, s). A) Mean (full bar), Median (dotted bar), and density distribution of MD for the six experimental conditions. The three speed conditions are annotated on the x-axis: Slow (blue), Spont (green) and Fast (red). The two directions are marked up within each bar: Down (D, dark color), Up (U, light color). Black lines and stars on the top of each histogram represent statistical differences. B) Mean (full bar), Median (dotted bar), and density distribution of MD inter-trial variability (MD CV) for the six experimental conditions.

For intra-subject variability (MD CV; Figure 2B), we observed strong inter-trial consistency within speed conditions (mean CV< 0.2 for the three speeds). The 3×2 ANOVA showed a main effect of speed (F(2,39)=8.39, p<0.01) and no main effect of direction (F(1, 40)=0.06, p=0.81). MD CV in Slow (0.16±0.06, p=0.69) was larger than that in Spont (0.15±0.05), larger than in Fast (0.12±0.08, p=0.016).

Interestingly, compared to a previous study (Gaveau & Papaxanthis, 2011) that assessed the temporal structure of vertical arm movements in a more restrictive situation (pointing at a salient target), our subjects chose to perform their movements at a generally slower speed, regardless of the speed condition (Gaveau’s data - Mean±STD - Fast:0.42±0.1; Natural:0.64±1.4; Slow:0.84±1.6). Interestingly, the intersubject variability associated with slow speed was similar between the two studies. Conversely, in the Spontaneous and Fast conditions, the inter-subject variability among our 40 subjects was lower than that observed by (Gaveau & Papaxanthis, 2011; 8 subjects). In particular, the large difference in the number of subjects between the two studies may have influenced the differences in the inter-subject variability values.

Finally, the correlation between the two directions (Up versus Down; Figure S1 in the Supplementary Material) showed that the MD was not significantly affected by the change in direction. Indeed, the MD adopted by a subject in one direction predicted MD in the other direction, independently of movement speed (r(39)>0.95; p<0.001 for the three speed conditions). This strong correlation, along with a regression line showing a slope close to 1 and an intercept close to 0 (slope = 0.8, intercept:0.22, slope = 0.85, intercept = 0.06, Spont: slope = 0.95, intercept = 0.01), shows that the effects of gravity on the motor system are generally anticipated and compensated to maintain the same movement duration.

### Maximum Velocity and Movement Vigor

#### Maximum Velocity

The 3×2 ANOVA ran on MaxV showed a main effect of speed (F(2,39)=202.55, p<0.0001) but not of movement direction (F(1,40)=0.82, p=0.37). MaxV significantly decreased by 37% and increased by 79% from Spont (1150 ± 303 mm.s^-1^) speed to Slow (721 ± 322 mm.s-1, p<0.001) and Fast (2060 ± 600 mm.s^-1^, p<0.001) conditions, respectively, and for the two directions (Figure 3A). The maximum velocity was strongly idiosyncratic, ranging from 629 to 1973 mm.s^-1^ at Spont speed.

**Figure 3.**
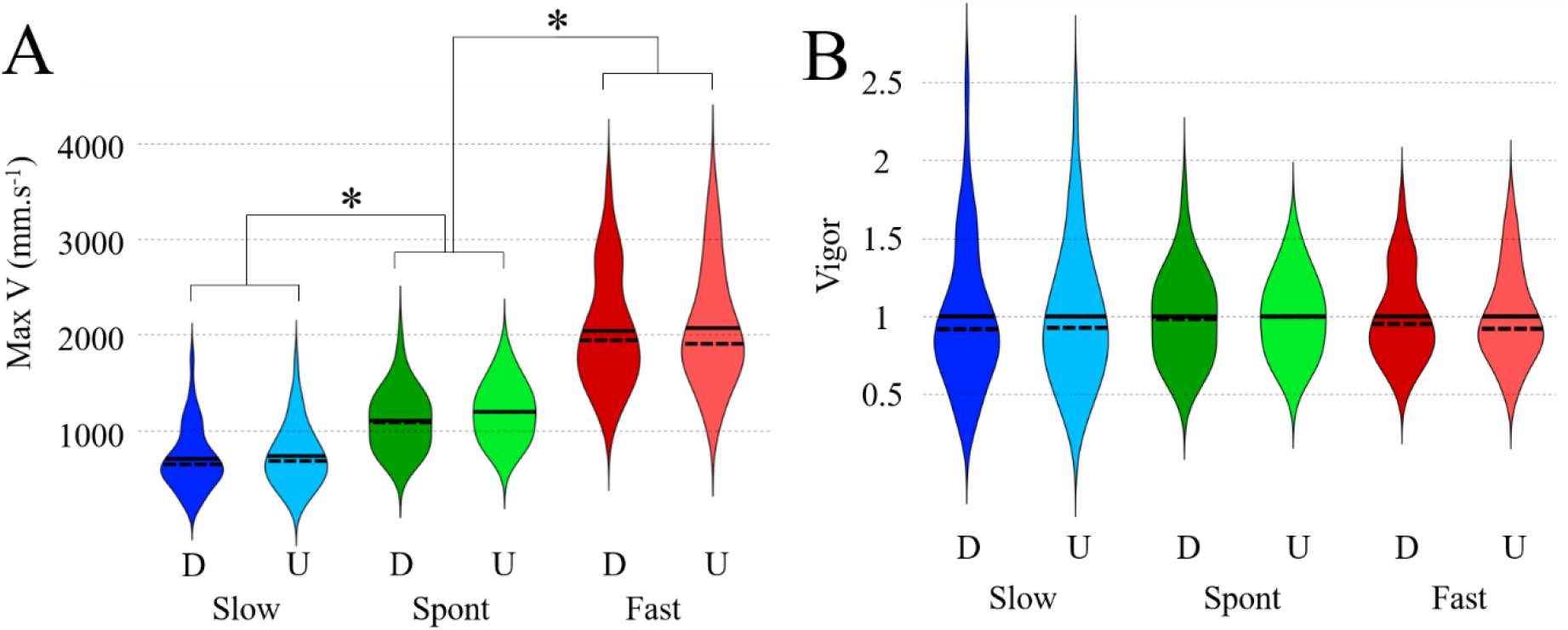
Effect of direction and speed on Maximum Velocity (Max V; in millimeters per second, mm.s^-1^) and Vigor. A) Mean (full bar), Median (dotted bar), and density distribution of A) MaxV and B) Vigor for the six experimental conditions. Same color code as in previous figures.

Finally, the correlation between the two directions (Up versus Down; Figure S2 in Supplementary Material) showed that MaxV was not significantly affected by the change in direction. Indeed, the MaxV adopted by a subject in one direction predicted the MaxV in the other direction, independently of the speed of the movements (r(39)>0.9; p<0.001 for the three speed conditions). These strong correlations are accompanied by regression lines with slopes close to 1 (slope = 0.96 for Slow; 6.98 for Spont; 1.00 for Fast). The intercepts were close to 0 in Slow and Fast but slightly superior in Spont (intercept = 55 for Slow; 1050 for Spont; 17 for Fast). Overall, these results suggest that MaxV was not significantly affected by the change in direction.

#### Movement Vigor

Movement vigor was introduced to further analyze the subjects’ Max V intersubject variability by comparing individual MaxV values to our sample mean. Movement vigor was strongly idiosyncratic (Figure 3B), ranging from 0.32 to 2.48 (1.00±0.45), from 0.57 to 1.78 (1.00±0.26) and from 0.52 to 1.76 (1.00±0.29) at Slow, Spont and Fast speed, respectively. Since vigor scores represent values that have been normalized by the same mean value, we did not perform an ANOVA analysis (mean comparison).

The Pearson tests performed between the two directions (Up versus Down; Figure S3 in Supplementary Material), showed that the movement Vigor adopted by a subject in one direction predicted the movement Vigor in the other direction, independently of the speed of the movements (r(39)>0.9; p<0.001 for the three speed conditions). Interestingly, when studying the associated regression lines, we observed slopes close to 1.00 and intercepts close to 0.00, particularly for Slow and Fast speed (slope=0.92, intercept=0.07 for Slow; slope=0.80, intercept=0.20 for Spont; slope=0.99, intercept=0.01 for Fast). Overall, these results confirm the stability of vigor across the two directions (i.e., robust, despite the mechanical perturbations induced by the change in direction).

### Spatial accuracy

#### Constant Error

The 3×2 ANOVA ran on CE (Figure 4A) showed a main effect of speed (F(2, 39)=4.73, p<0.01) and movement direction (F(1, 40)=47.11, p<0.0001), with no significant interaction (F(2, 39)=0.13, p=0.88). Post-hoc tests revealed a significant difference between Spont and Fast speed (p<0.01), but not between Spont and Slow (p=0.61) or Fast and Slow speeds (p=0.10). In particular, the Up and Down movements were respectively undershooting and overshooting in the Spont and Slow speed conditions (Up Spont: −5.35± 15; Down Spont: 13.05 ± 21; Up Slow:-2.08±16.29, Down Slow:16.38±18, Up Fast:3.50±10.72, Down Fast:25.00±27).

**Figure 4.**
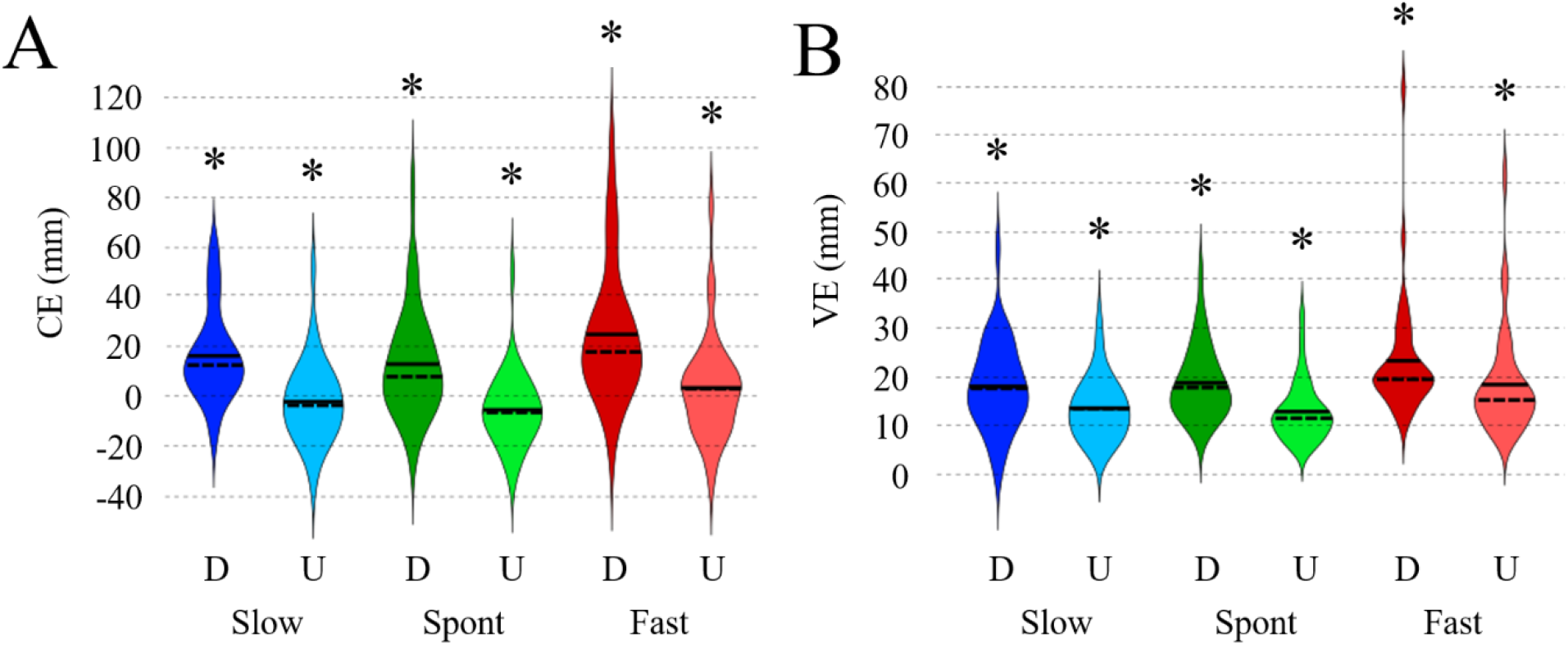
Effect of direction and speed on Constant and Variable Error. Mean (full bar), Median (dotted bar), and density distribution of A) the constant error (CE, in mm), and B) the variable error (VE, in mm), for the six experimental conditions. Same color code as in previous figures.

The Pearson tests performed between the two directions (Up versus Down; Figure S4 in Supplementary Material) showed a significant correlation between Fast Down CE and Fast Up CE (R=0.52, p<0.001), but no correlation in Spont and Slow conditions (Spont: R=0.20, p=0.24; Slow: R=0.30, p=0.09). Furthermore, regardless of the speed condition, the trends line associated to these correlations had slopes largely inferior to 1 and intercepts greater than 0 (slope=0.25, intercept=-6.12 for Slow; slope=0.14, intercept=-7.15 for Spont; slope=0.40, intercept=-6.45 for Fast). These results indicate that changes in direction affected CE differently across the subjects.

#### Variable Error

The 3×2 ANOVA ran on VE (Figure 4B), showed a main effect of speed (F(2, 39)=7.38, p<0.001) and movement direction (F(1, 40)=16.54, p<0.0001), without significant interaction (F(2, 39)=0.11, p=0.89).

The Pearson tests performed between the two directions (Up versus Down; Figure S5 in the Supplementary Material) showed significant correlations at the three experimental speeds (Spont: R=0.74, p<0.001; Slow: R=0.46, p<0.05; Fast: R=0.55, p<0.001). Furthermore, regardless of the speed condition, the trends line associated to these correlations had slopes largely inferior to 1 and intercepts greater than 0 (slope=0.37, intercept=6.31 for Slow; slope=0.61, intercept=1.36 for Spont; slope=0.49, intercept=7.11 for Fast). These results indicate that changes in direction affected CE differently across the subjects. VE, as CE, is affected differently across the subjects by changes in direction.

### Movement Timing

In agreement with previous results (Berret, Gauthier, et al., 2008; Gaveau et al., 2016; Papaxanthis et al., 1998) the acceleration time was, on average, less than 50% of the total duration (i.e., TPV < 0.5) for upward pointing and close to 50 % (that is, TPV ≈0.5) for downward pointing (Figure 5A and Table S1 for the mean values). The 3×2 ANOVA ran on TPV revealed a main effect of speed (F(2, 39)=9.22, p<0.001) and movement direction (F(1, 40)=75.29, p<0.0001) without significant interaction (F(2, 39)=0.54, p=0.58). Indeed, mean TPV across the two experimental directions significantly decreased from Down (0.49 ± 0.06) to Up (0.43 ± 0.05) direction (Figure 5A). TPV was not different from Fast (0.48 ± 0.05) to Spont (0.47 ± 0.05, p=1) and significantly decreased from Spont to Slow speed (0.44 ± 0.07, p<0.01).

**Figure 5.**
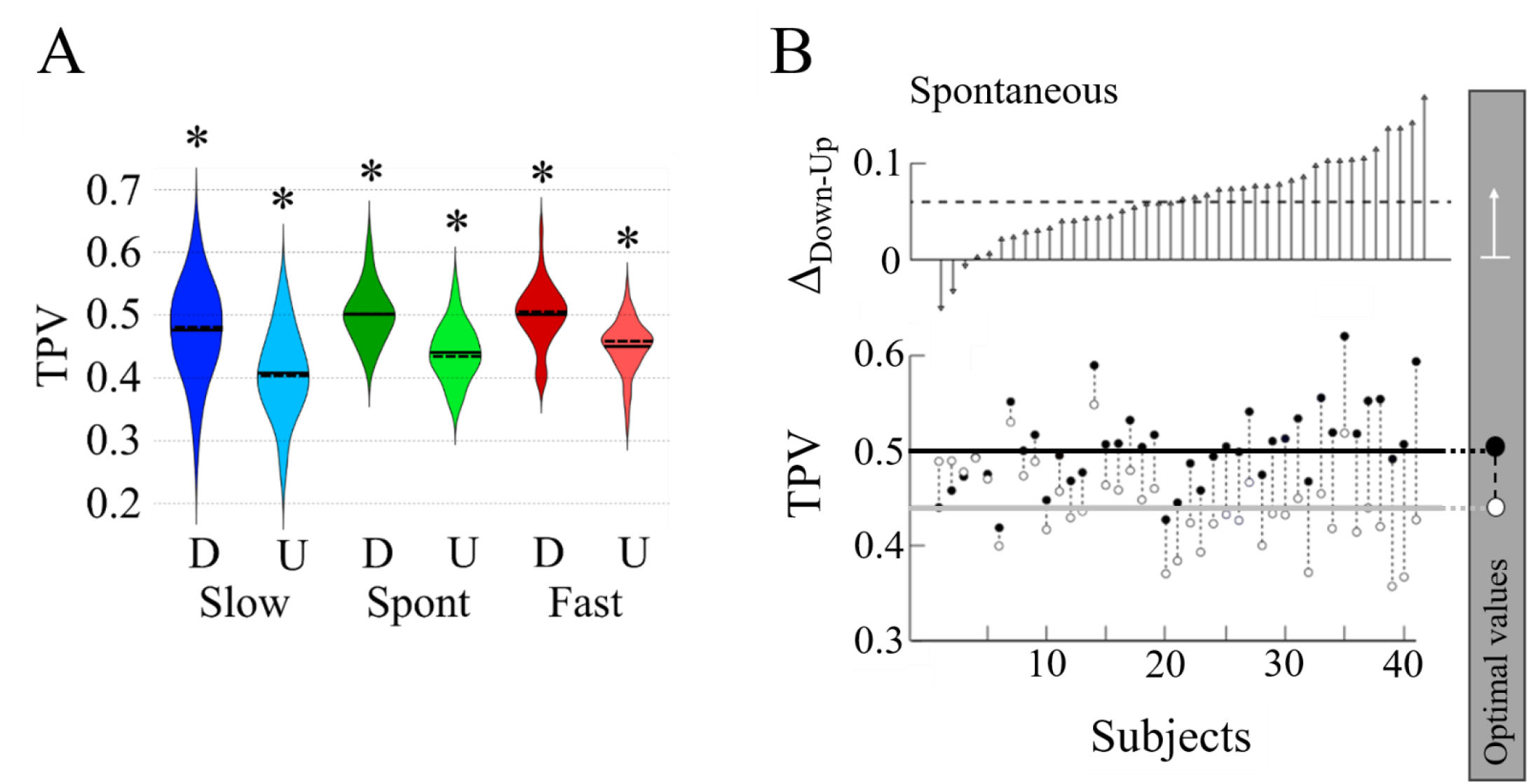
Time to Peak Velocity. A) Mean (full bar), Median (dotted bar), and density distribution of TPV for the six experimental conditions. Same color code than in previous figures. B) High panel: individual TPV Delta values (Down – Up; arranged in ascending order), for the Spontaneous speed condition. Low panel: associated Down (black dot) and Up (white dot) TPV values. Optimal values predicted by the smooth-effort model (Gaveau et al., 2016) are illustrated within the right grey panel.

Interestingly, across the 41 participants tested TPV values in the Down direction ranged from 0.40 to 0.64 (0.50±0.05) for Fast speed, from 0.42 to 0.62 (0.50±0.04) for Spont speed, and from 0.28 to 0.62 (0.48±0.07) for Slow speed. In the Up direction, TPV values ranged from 0.33 to 0.54 (0.45±0.04) for Fast speed, from 0.36 to 0.55 (0.44±0.04) for Spont speed, and from 0.26 to 0.55 (0.41±0.07) for Slow speed. These data show that the variability associated with fast and spontaneous speeds are comparable regardless of the direction of movement. In contrast, in the slow-speed condition, the variability and amplitude of the range increased.

The correlations performed between the two directions (Up versus Down; Figure S6 in the Supplementary Material) showed that the Down TPV predicted the Up TPV. Indeed, participants who spent longer (or shorter) time accelerating downward also spent longer (or shorter) time accelerating upward (r(39)=0.68, p<0.001 for Fast; r(39)=0.48, p<0.01, for Spont; r(39)=0.48, p<0.001 for Slow). However, regardless of the speed condition, the trend line associated with these correlations had slopes largely inferior to 1 and intercepts different than 0 (slope=0.43, intercept=0.20 for Slow, slope=0.47, intercept=0.20 for Spont; slope=0.58, intercept=0.16 for Fast). These results indicate that changes in direction affected the TPV differently across the subjects.

To further evaluate the influence of the directions on the TPV, we evaluated the delta TPV, defined as the difference between Down and Up TPV. This parameter reflects the presence of vertical TPV asymmetry and has been previously described as an efficient estimator of energetic optimization (Gaveau et al., 2016). In the Spont condition (Figure 5B), delta TPV varied from −0.05 to 0.17 (0.06±0.04). Approximately seven participants (on 41; Figure 6) reached the optimal value of delta predicted by the smooth effort model (Gaveau et al., 2016). Similarly for Slow and Fast speed (Figure 6), delta TPV varied from −0.17 to 0.18 (0.07±0.07) and from −0.06 to 0.11 (0.05±0.04), respectively. Approximately six participants for Fast and eight participants for Slow reached the optimal value of delta. These observations showed that despite the mean delta TPV values that are consistent with the prediction of the smooth effort model, the examination of individual values showed that most of the subjects were out of this optimal range. Our data seem to point toward a continuum between subjects adopting a certain mode of behavior favoring small differences across down and up TPV to subjects adopting another mode of behavior producing large differences across down and up TPV.

**Figure 6.**
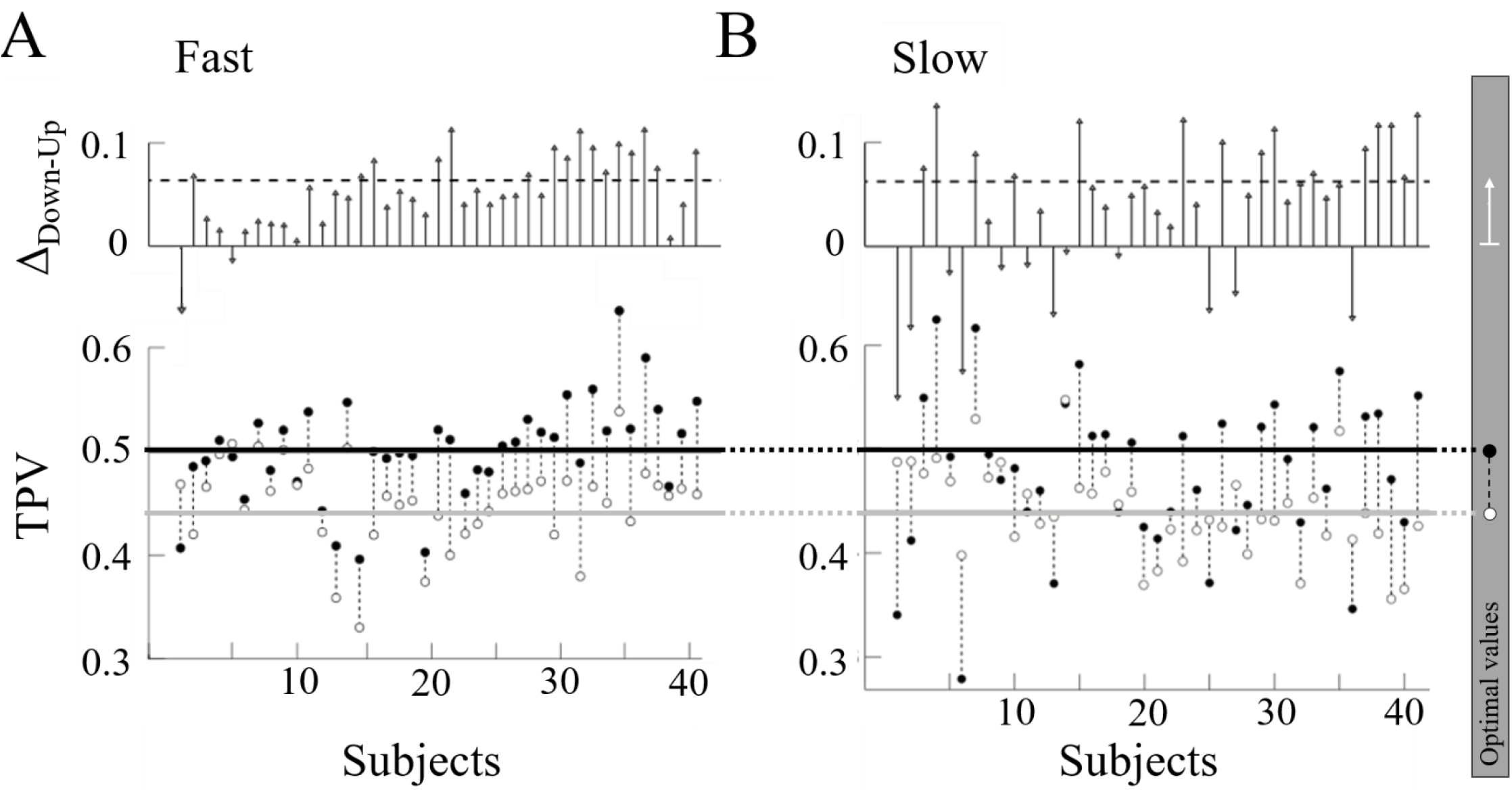
Time to Peak Velocity. High panels represent the individual TPV Delta values (Down – Up; arranged in the same order as Figure 5B), for the A) Fast and B) Slow speed conditions. Low panels show the associated Down (black dot) and Up (white dot) TPV values. Optimal values predicted by the smooth-effort model (Gaveau et al., 2016) are illustrated within the right grey panel.

### Net Metabolic Rate

To evaluate whether this variability may still be compatible with the minimization of energy expenditure, we computed the Net Metabolic Rate (NMR) for each subject and each condition Since subject-wise Up NMR values are negative and Down and Up NMR values are comparable (Figures S8 in Supplementary Material), we thus compared delta TPV with the individual mean of absolute NMR values 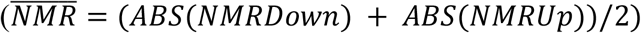. To do so, we compared the individual delta TPV values with their associated 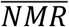 for each experimental condition (Figure 7 for Spont, and Figure S7 for Slow and Fast). The Pearson test showed a very weak correlation between 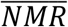 and delta TPV, regardless of the condition (Spont: R=0.09, p=0.67; Fast: R=-0.20, p=0.36; Slow: R =-0.13, p = 0.52). Finally, to compare our NMR computation with the optimization of the smooth effort criterion (Gaveau et al., 2016), we looked at the NMR scores of subjects having a Δ score close to an optimal value (0.3≤Δ≤0.8) and subjects having a Δ score far from an optimal value (Δ<0.3 & Δ>0.8; Table 1). Indeed, for all conditions, we observed smaller value of NMR for the subject having a Δ score closer to optimal values. These results suggest that the large variability observed across the individual delta TPV cannot be explained by the minimization of energy expenditure, as represented by our NMR computation.

**Figure 7.**
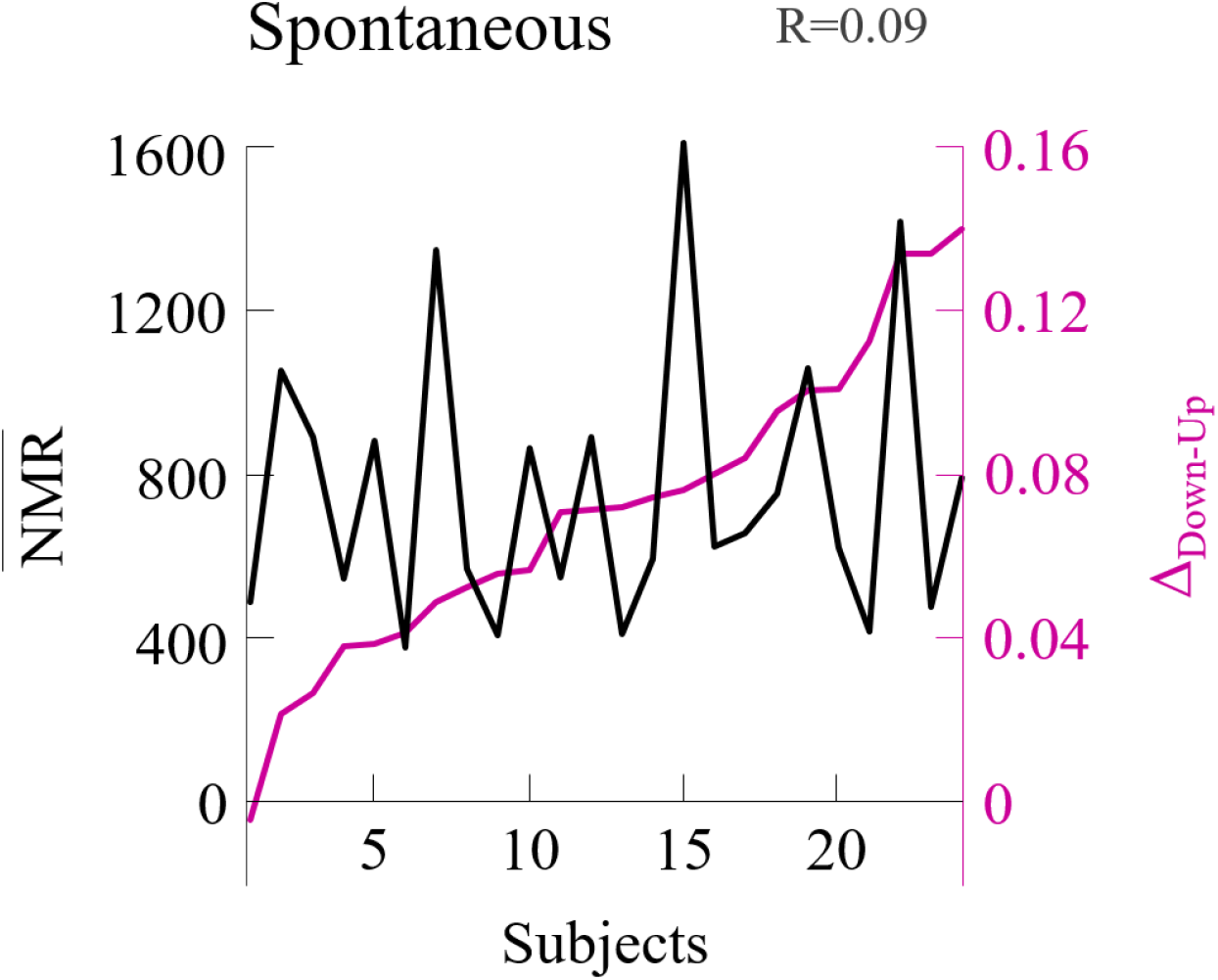
Comparison of individual TPV Delta (Δ, Down – Up) and Net Metabolic Rate 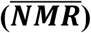. Distribution of subjects TPV Delta (Δ_Down-Up_, Red curve arranged in ascending order) and associated individual Net Metabolic Rate (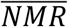, Black curve) for the Spontaneous speed condition.

**Table 1.** Net Metabolic Rate values according to Δ values group. Mean and standard deviation of NMR values for the three speed and the two direction conditions, for subjects having Δ around smooth-effort-optimal values (between 0.3 and 0.8, first row) and for subjects with Δ outside this optimal range (Δ<0.3 or Δ>0.8, second row). The number of subjects (N) in each category is written in grey at the bottom of each cell.

|  | **Down**<br>**Slow** | **Down**<br>**Spont** | **Down**<br>**Fast** | **Up**<br>**Slow** | **Up**<br>**Spont** | **Up**<br>**Fast** |
| --- | --- | --- | --- | --- | --- | --- |
| **$0.3 \leq \Delta \leq 0.8$** | 535.09<br>$\pm 249$<br>*N=9* | 739.89<br>$\pm 377$<br>*N=13* | 1128.71<br>$\pm 428$<br>*N=12* | -487.42<br>$\pm 206$<br>*N=9* | -745.30<br>$\pm 373$<br>*N=13* | -1128.85<br>$\pm 419$<br>*N=12* |
| **$\Delta < 0.3$**<br>**or**<br>**$\Delta > 0.8$** | 585.75<br>$\pm 355$<br>*N=15* | 785.57<br>$\pm 313$<br>*N=11* | 1163.80<br>$\pm 368$<br>*N=12* | -552.71<br>$\pm 362$<br>*N=15* | -781.16<br>$\pm 301$<br>*N=11* | -1154.91<br>$\pm 377$<br>*N=12* |

## Discussion

This study aimed to verify the existence of inter-individual differences during arm pointing in the vertical plane, a task that is highly dependent on the mechanical effect of gravity. Based on the idea that a pointing task performed under conditions of maximum speed towards a salient target can flatten the distribution of individual responses, we used a pointing task relaxing external constraints (i.e. with limited temporal and spatial constraints). We found a significant variability of individual modes of interaction with the gravitational force field and idiosyncratic motor responses. The analysis of energy costs revealed the presence of suboptimal motor response suggesting that arm movement timing is not exclusively defined by the energy criterion. In the following paragraphs, these results are discussed considering the role of subjective values in guiding individual motor planning achieved with or against the force of gravity.

### Movement duration and vigor are idiosyncratic and invariant along up and down directions

MD was found to be invariant along the two directions for all three-speed conditions (slow, spontaneous, and fast). However, among all participants tested, individual MD was idiosyncratic, with some participants consistently performing the task for three times the duration of others. Further, individual MD was predictable from the values recorded at spontaneous speed and the slowest/fastest participants at spontaneous speed remained the slowest/fastest at slow and fast speeds, highlighting the stability of each pace. This observation is unexpected, particularly at slow speed since it is well-known that humans tend to avoid unnatural slow speeds. Indeed, movements with a longer duration require more neural and attentional resources (Guigon et al., 2019; Park et al., 2017) and are judged more effortful than shorter movements against the same force (Van Der Wel et al., 2009). As a result, we should have recorded a reluctance to slow down the movement (since movement durations were not imposed), resulting in a distribution of MD values at slow speed close to the duration recorded at spontaneous speed.

From a decision-making perspective, movement duration has been associated with different motivations (Cisek, 2005; Cisek & Kalaska, 2010; Wolpert & Landy, 2012). Fast individuals reduce the cost of time by minimizing the implicit delay in reward acquisition but simultaneously increase the metabolic cost of the motor response, and vice versa for a long MD (Shadmehr et al., 2016). The wide range of durations recorded here at slow speed could reflect an adaptation process during actions performed at unusual speed, guided by individual internal values minimizing either effort or time, as demonstrated for pointing movements in the horizontal plane (Berret & Jean, 2016).

Furthermore, unlike horizontal orientation, upward and downward movements are associated with an opposite sense of effort. For movements of similar amplitude and duration, the movement performed against gravity is estimated as more effortful (Proske, 2005, 2006; Van Beek et al., 2013). Here, such asymmetric “mental effort” has not been confirmed at the behavioral level, where MD is currently direction-independent. Overall, i) the ability of participants to easily adopt speeds that deviate from their spontaneous speed, ii) the idiosyncrasy of MD, and iii) the independence of MD from pointing direction suggest that the minimization of energy is not a constraint that can be generalized to all participants.

Similarly, when participants were required to point at an unusually slow speed, individual vigor was predictable from the spontaneous speed values. Again, individual vigor scores were not affected by movement direction and were idiosyncratic, especially at slow and spontaneous speeds. At high speed, the distribution of individual vigor values became more compact, and the spread between participants decreased. The invariance of vigor across different sensorimotor contexts (i.e., with or against the force of gravity, changes in inertia induced at fast speeds, and the neural adjustments required when pointing at an unusually slow speed) indicate that this variable is an abstract component of the task goal rather than a representation of its mechanical details (Bartlett, 1932; Bernstein, 1967; Schmidt, 1975).

Furthermore, the identification of vigorous and slow participants suggests that this kinematic feature is a subjective value guiding task execution. Subjective value here refers to a relatively stable and distinctive attribute of individuality (Labaune et al., 2020). Accordingly, several approaches propose that movement vigor is the motor outcome of a neural mechanism related to the subjective estimation of time and effort (Shadmehr et al., 2019). The individualization of this variable as well as its conservation across different mechanical and sensory situations is consistent with the idea that vigor reveals a personality trait. For example, it tends to corroborate a psychological trait relating to the propensity for boredom and the cost of time (Berret et al., 2018). In our case, participants with high movement vigor might reveal expensive and impatient participants for whom time determines proximity to reward. Conversely, slow pointing would depict participants who “invest” more in minimizing effort. The diversity of individual vigor across the different modalities tested here could therefore reflect motor stylistic variations during a simple pointing task. Concerning the neural substrates involved, the dorsal striatum could be part of the mechanism responsible for modulating vigor in response to changes in reward predictability (Jurado-Parras et al., 2020; Opris et al., 2011).

### Movement timing and minimization of energy cost

In agreement with previous studies (Berret, Gauthier, et al., 2008; Gaveau et al., 2016; Papaxanthis et al., 1998), upward movement timing was found asymmetric (i.e., with shorter normalized time to accelerate than to decelerate). Direction-dependent kinematic of movement performed in the vertical plane has been identified as putative exploitation of gravity force field to initiate downward movement and slow upward movement (Pozzo et al., 1998). Asymmetric movement kinematic has been formalized in a minimum smooth-effort model (Berret, Gauthier, et al., 2008) that would reflect an optimal integration of gravity force field minimizing the motion effort (Gaveau et al., 2016, 2021).

Following the energy minimization assumption, individual TPV values were expected to cluster around 0.52 and 0.42 for the downward and upward directions, respectively. On this basis, the TPV values recorded at fast and spontaneous speeds are partially at odds with the idea that all participants intended to minimize energy cost by taking advantage of a generic internal model of gravity. 48% of the 41 participants tested performed suboptimal downward movement (with a longer deceleration duration compared to acceleration duration) at spontaneous and fast speed. A movement timing dedicated to economical pointing predicts that the upward and downward kinematics are adapted to assistive and resistive effects of gravity. Our results did not show such a pattern. Conversely, the downward TPV values are positively correlated with the upward TPV values, so when the timing of the movement is optimal in one direction, it is detrimental to the minimization of energy in the other direction. Thus, for a given direction, the TPVs vary from subject to subject so that the Delta TPVs (TPV down - TPV up) are relatively high in some participants but relatively low in others.

While providing a clear query about the role of energy minimization in vertical arm pointing motion planning, it should nevertheless be noted that we assessed energy using a Net Metabolic Rate formula (Shadmehr et al., 2016). Particularly, this metabolic cost takes into account physiological and kinematic variables without considering parameters known to be related to energy expenditure such as oxygen consumption or EMG patterns.

### Conclusion

The present results showed the existence of idiosyncratic kinematic characteristics that are consistent for various velocities, mechanical modalities (upward or downward), and gravity contexts. This indicates that individual components determine the kinematics of the action performed in the vertical plane. Indeed, the observation of a continuum of behaviors ranging from fast to slow participants is not consistent with the idea that movements performed in the vertical plane rely exclusively on a generic internal model of gravity minimizing energy expenditure. Rather, our data demonstrate that individual-specific internal values (and the subjective rewards associated with each of these values) influence the kinematic characteristics of arm movements in the vertical plane. However, humans perform remarkably well in endurance races that encourage the minimization of mechanical effort, as demonstrated by the diverse evolutionary picture of bipedalism (Bramble & Lieberman, 2004; Carrier, 2011). Therefore, we hypothesize that energy minimization, resulting from the evolutionary advantage it provides, combines with individual internal values formed during ontogeny for action planning. Further investigations should describe the relative weight of these personal traits with basic constraints in individual motor control.

