## Supplementary Material for "Evidence for subjective values guiding cognitive integration of gravity in a free speed manifold pointing task"

| | **Down Slow** | **Down Spont** | **Down Fast** | **Up Slow** | **Up Spont** | **Up Fast** |
|---|---|---|---|---|---|---|
| **MD (s)** | 1,75 *±0,12* | 1.04 *±0,04* | 0,60 *±0,03* | 1,63 *±0,10* | 0,94 *±0,04* | 0,57 *±0,03* |
| **Max V ($mm.s^{-1}$)** | 701 *±51* | 1107 *±48* | 2049 *±92* | 728 *±50* | 1196 *±45* | 2066 *±95* |
| **TPV** | 0,48 *±0,01* | 0,50 *±0,01* | 0,50 *±0,01* | 0,41 *±0,01* | 0,44 *±0,01* | 0,45 *±0,01* |
| **Cst Error (mm)** | 16.38 *±2.89* | 13.05 *±3.40* | 25.00 *±2.89* | -2.08 *±2.31* | -5.35 *±2.30* | 3.50 *±2.30* |

**Table S1 – Main motor variables.** Mean, and Standard Error of the main movement parameters: movement duration (MD, in seconds), maximum velocity (Max V, in $mm.s^{-1}$), time to peak velocity (TPV), and constant error (CE, in millimeters), for the 6 experimental conditions (Downward-Slow, Downward -Spontaneous, Downward -Fast, Upward-Slow, Upward - Spontaneous, Upward -Fast).

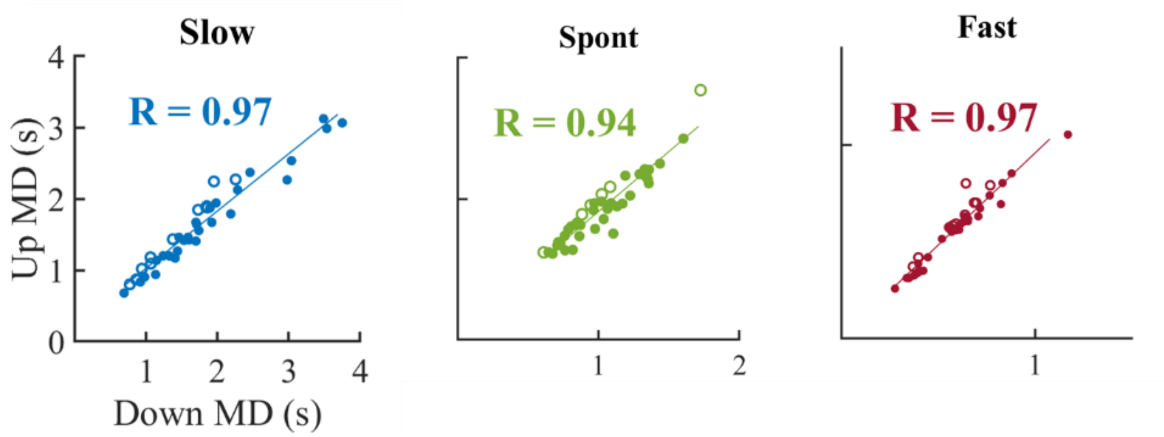


**Figure S1.** Scatter plots of MD comparing movement direction (x-axis: Down, y-axis: Up) across subjects, in the three speed conditions (Slow: blue, Spont: green and Fast: red). Within each color, white dots indicate subjects having higher MD in the Up direction (compared to Down), and vice versa for dark dots.

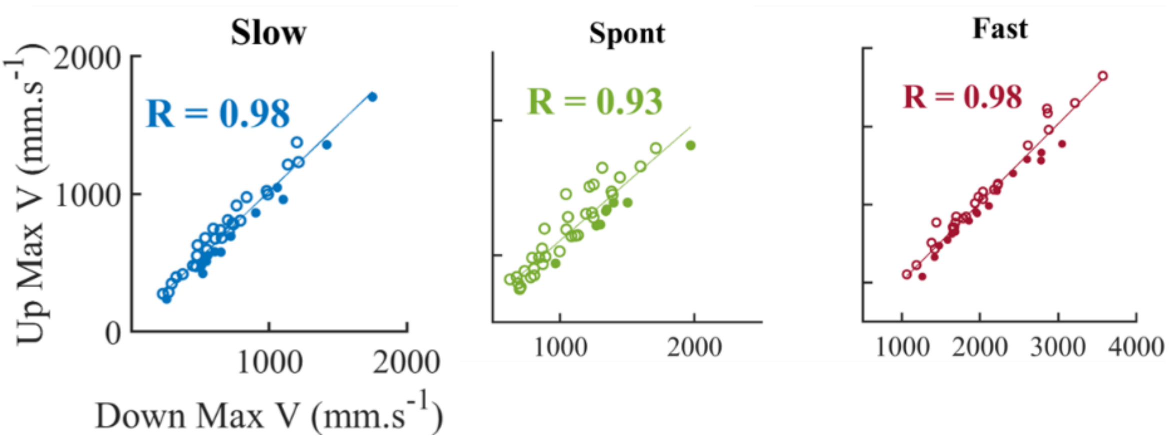


**Figure S2.** Scatter plots of MaxV comparing movement direction (x-axis: Down, y-axis: Up) across subjects, in the three speed conditions. Same color code as in previous figures.

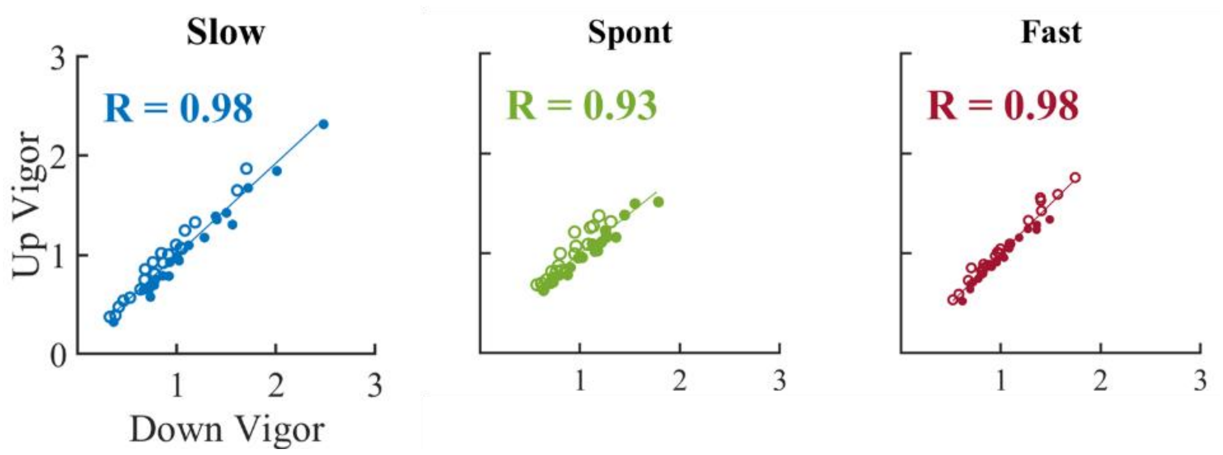


**Figure S3.** Scatter plots of movement Vigor comparing movement direction (x-axis: Down, y-axis: Up) across subjects, in the three speed conditions. Same color code as in previous figures.

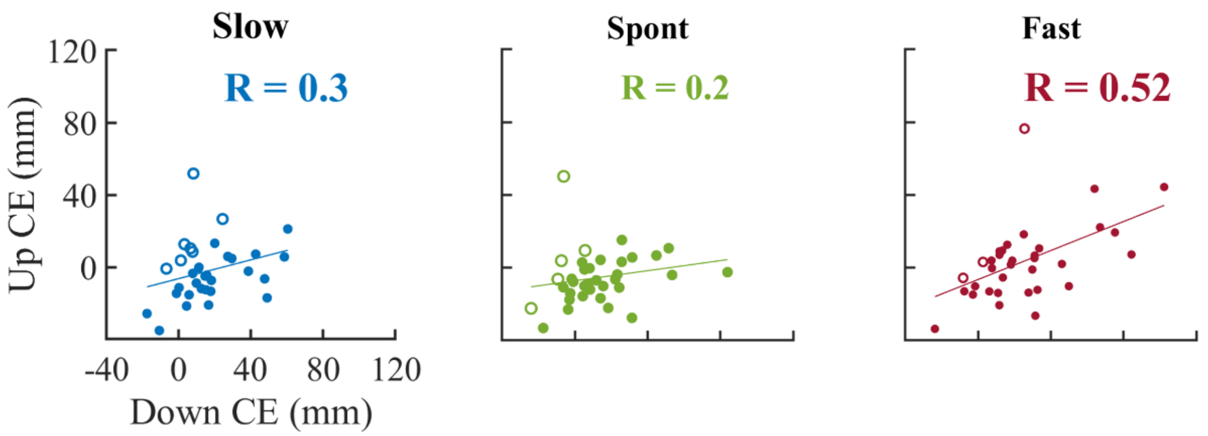


**Figure S4.** Scatter plots of CE comparing movement direction (x-axis: Down, y-axis: Up) across subjects, in the three speed conditions. Same color code as in previous figures.

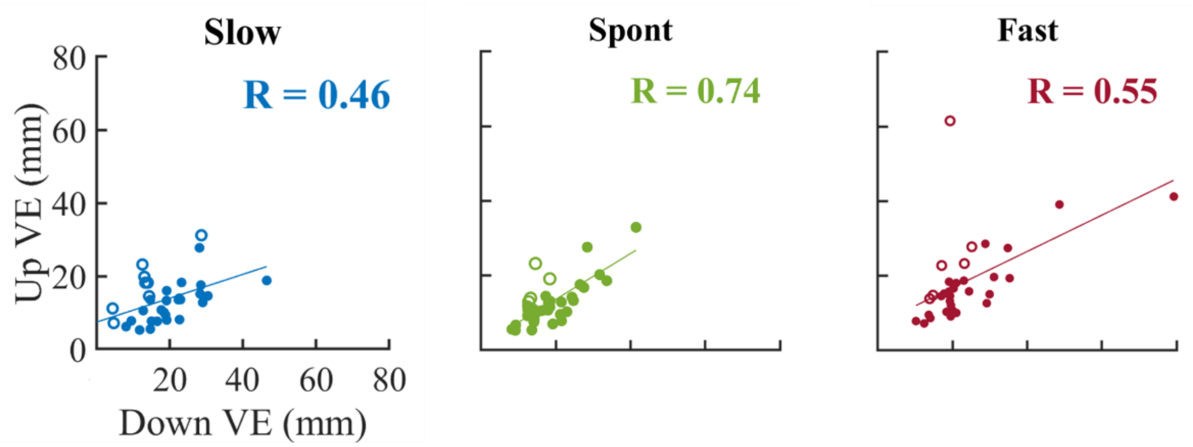


**Figure S5.** Scatter plots of VE comparing movement direction (x-axis: Down, y-axis: Up) across subjects, in the three speed conditions. Same color code as in previous figures.

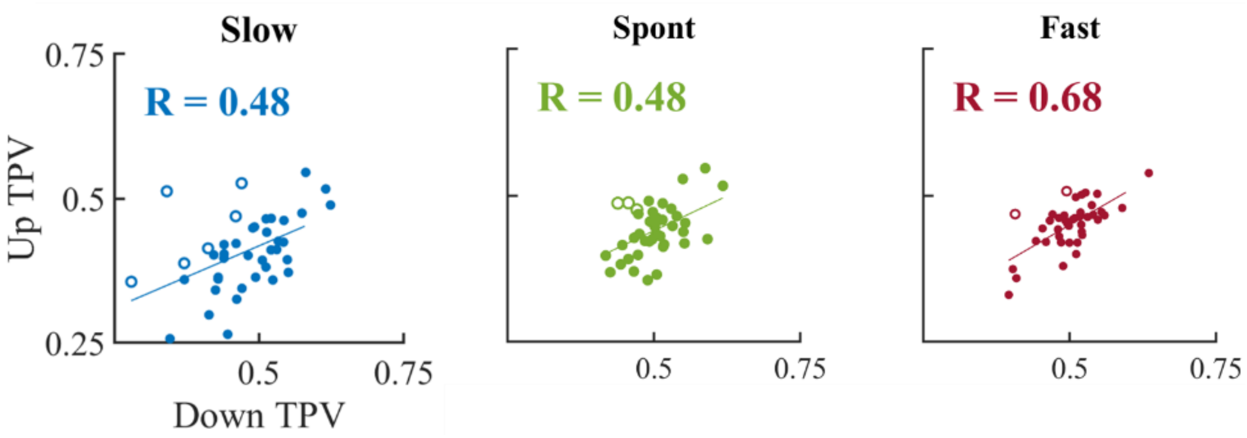


**Figure S6.** Scatter plots of TPV comparing movement direction (x-axis: Down, y-axis: Up) across subjects, in the three speed conditions. Same color code as in previous figures.

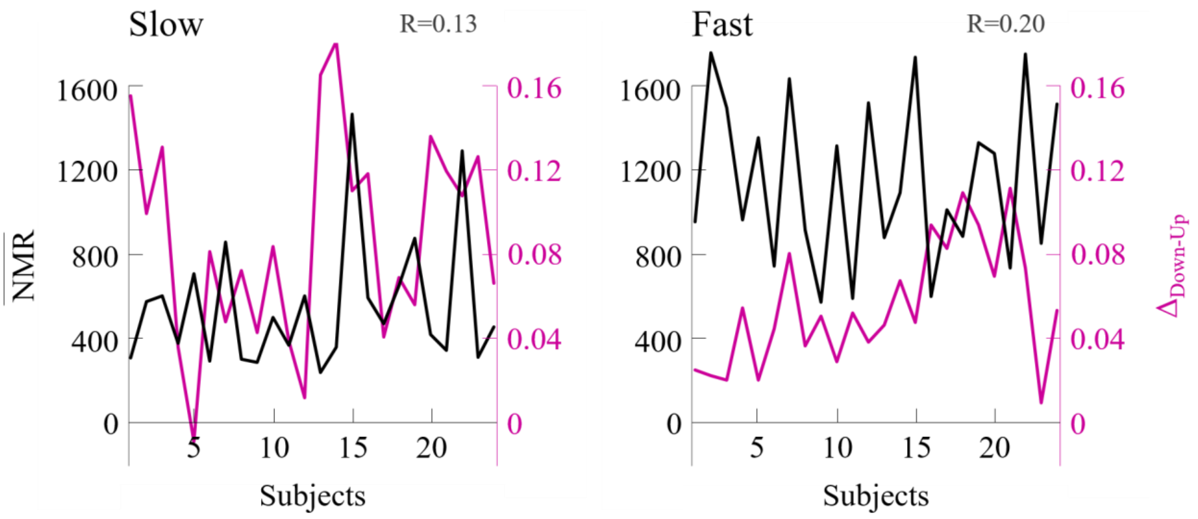


**Figure S7. Comparison of individual TPV Delta (Δ, Down – Up) and Net Metabolic Rate ($\overline{\boldsymbol{NMR}}$).** Distribution of subjects TPV Delta ($\Delta_{\text{Down-Up}}$) and associated individual Net Metabolic Rate ($\overline{NMR}$, black curve) for A) the Slow and B) the Fast speed condition.

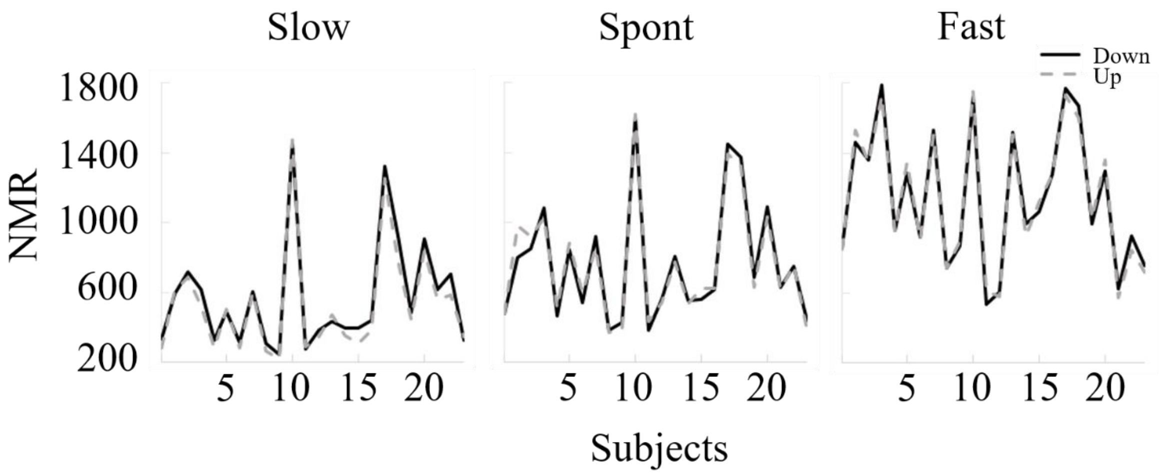


**Figure S8.** Comparison of Downward (black curve) and Upward (dotted grey curve) Net Metabolic Rate (NMR), for a typical subject and for the three speed conditions (Slow - left, Spont – middle, Fast - right).
